# Autophagy disruption primes CAR-T cell metabolism for sustained rejection of ovarian tumors

**DOI:** 10.1101/2025.10.09.681473

**Authors:** Gillian A Carleton, Sébastien Levesque, Lauren G Zacharias, Joao San Patricio, Tracey Sutcliffe, Peter H Watson, Ralph J DeBerardinis, Yannick Doyon, Julian J Lum

**Affiliations:** Department of Biochemistry and Microbiology, University of Victoria, Victoria, BC; Trev and Joyce Deeley Research Centre, BC Cancer, Victoria, BC; Centre Hospitalier Universitaire de Québec Research Center – Université Laval, Québec, QC; Université Laval Cancer Research Centre, Québec, QC; Present Address: Division of Hematology/Oncology, Boston Children’s Hospital. Department of Pediatric Oncology, Dana-Farber Cancer Institute. Harvard Stem Cell Institute. Broad Institute of MIT and Harvard. Department of Pediatrics, Harvard Medical School, Boston, MA; Children’s Research Institute, UT Southwestern, Dallas, TX; Animal Care Services, University of Victoria, Victoria, BC; Biobanking and Biospecimen Research Services, Deeley Research Centre, BC Cancer, Victoria, BC; 9Howard Hughes Medical Institute, UT Southwestern Medical Center, Dallas, TX; Eugene McDermott Center for Human Growth and Development, UT Southwestern Medical Center, Dallas, TX

## Abstract

T-cell based immunotherapies such as chimeric antigen receptor T (CAR-T) cell therapy face substantial hurdles when confronting solid tumors such as ovarian cancer, where metabolic constraints in the tumor microenvironment limit T cell infiltration and function. In particular, T cells exposed to nutrient deprivation and hypoxia upregulate autophagy, a lysosomal degradation pathway that negatively regulates effector responses. Here, we used CRISPR-Cas9 to target a folate receptor alpha (αFR) CAR expression cassette into the locus of the essential autophagy gene *ATG5*, thereby generating autophagy-deficient CAR-T cells in a single editing step.

Targeted metabolite profiling revealed that deletion of *ATG5* induced widespread metabolic reprogramming characterized by increased glucose and amino acid uptake. Functionally, *ATG5*-knockout CAR-T cells maintained high cytolytic activity when assayed in patient-derived ascites *in vitro*, and exhibited superior and long-lasting tumor control against ovarian tumors *in vivo*.

Taken together, our results suggest that deletion of *ATG5* metabolically primes CAR-T cells for enhanced cytotoxicity in immune-suppressive conditions, thereby improving the therapeutic potential of αFR CAR-T cells for ovarian cancer immunotherapy.

## INTRODUCTION

Chimeric antigen receptor T (CAR-T) cell therapy has transformed the standard of care for the treatment of cancer. However, the effectiveness of CAR-T therapy has been largely restricted to hematological malignancies (1), and similar results have yet to be achieved in cancers that form solid tumors, such as ovarian cancer (2). Ovarian cancer remains a clinical challenge, marked by significant genomic heterogeneity, high metastatic potential, and recurrence rates exceeding 70% (3,4). In high-grade serous ovarian carcinoma (HGSOC), the most prevalent and lethal subtype, five-year survival rates are poor, at just 41% and 20% for patients with stage III and stage IV disease, respectively (5). Despite major advances, the promise of immunotherapy has yet to translate into clinical benefit for ovarian cancer patients. This is especially apparent in the case of CAR-T therapy, as multiple Phase 1 CAR-T trials targeting the ovarian-associated antigens mesothelin or folate receptor alpha (αFR) have failed to achieve significant clinical responses (6–10). A similar lack of efficacy has been observed with immune checkpoint inhibitors such as nivolumab and pembrolizumab, which suggests that ovarian cancer may be broadly resistant to immunotherapeutic strategies (11). Emerging evidence points to the immunosuppressive and metabolically-restricted tumor microenvironment (TME) as a key barrier to effective immune engagement and CAR-T cell activity in this disease (12).

In advanced ovarian cancer, malignant cells originating from the fallopian tube frequently disseminate to sites within the peritoneal cavity, including the omentum, mesentery, and liver (13). This pattern of peritoneal metastasis is often accompanied by the accumulation of malignant ascites, a pro-inflammatory fluid that contributes to disease progression (14). For patients with advanced ovarian cancer, the TME therefore comprises both solid and liquid compartments, each containing a heterogenous population of malignant, stromal, and suppressive immune cells capable of inhibiting T cell function (15). Among the various immunosuppressive mechanisms within the TME, metabolic dysfunction, driven by both nutrient restriction and accumulation of immune-inhibitory metabolites, is a consistent and prominent feature across both compartments. In the solid TME, tumor cells and cancer-associated fibroblasts (CAFs) consume essential nutrients such as glucose and L-arginine (16–18), and produce elevated levels of the inhibitory metabolite lactate, which has been shown to impair cytokine production by T cells (19,20). In the ascites, increased amino acid uptake and catabolism exhibited by disseminated tumor cells and CAFs leads to depletion of phenylalanine, tyrosine, and tryptophan, along with production of inhibitory downstream metabolites such as kynurenine (21,22). Lastly, hypoxia further compounds metabolite stress in both environments (23–25).

Hypoxia and amino acid deprivation, hallmarks of the ovarian TME, are potent activators of autophagy, a catabolic pathway that recycles intracellular components through lysosomal degradation (26,27). While autophagy is essential for cellular homeostasis, mounting evidence indicates that activation of this pathway attenuates T cell effector function (28,29), likely through the preferential degradation of molecules present in cytoplasmic vesicles, such as granzymes, perforin, and nutrient transporters (30). However, the immunosuppressive effects of autophagy on T cell function can be reversed through targeted disruption of core autophagy genes (31).

Here, we employed CRISPR-Cas9 to insert an αFR-targeting CAR expression cassette directly into the locus of the essential autophagy gene *ATG5*, an “all-in-one” editing approach that generates autophagy-deficient CAR-T cells in a single editing event (hereafter referred to as “*ATG5*-knockout CAR-T cells”). Targeted metabolite profiling revealed that deletion of *ATG5* increased glucose and amino acid uptake, thereby priming *ATG5*-knockout CAR-T cells for increased cytolytic capacity. Accordingly, *ATG5*-knockout CAR-T cells exhibited superior and long-lasting control of OVCAR3 ovarian tumors *in vivo*. These findings demonstrate the therapeutic potential of autophagy disruption in CAR-T cells as a strategy to overcome metabolic suppression in the ovarian TME, and identify metabolic reprogramming as a key factor linking autophagy disruption to enhanced effector function. This approach provides a blueprint for engineering next-generation CAR-T cells with optimized metabolic capacity for ovarian cancer immunotherapy.

## RESULTS

### Design and validation of a strategy for single-step editing at *ATG5*

First, we developed a gene-editing strategy for generating autophagy-deficient CAR-T cells in a single editing event. We used the UCSC Genome Browser (32) to determine the first intron shared by all *ATG5* variants, which corresponded to intron 2 of the canonical *ATG5* sequence (transcript variant 1). Of note, we selected an intron rather than an exon to ensure that autophagy inactivation would occur only in cells with successful donor integration. After examining the sequence for intron 2, we identified three regions free of repetitive DNA elements suitable for editing (Supplementary Figure 1A). We designed gRNAs using CRISPOR (33), and screened them for on-target activity using a mismatch-sensitive assay. gRNA screening identified a highly active gRNA (*ATG5*-I2A-824) that was selected for all future experiments (Supplementary Figure 1B). After gRNA selection, we transfected K562 cells with a donor cassette encoding a fluorescent reporter gene, mScarlet-I (Supplementary Figure 2A). As expected, clones with successful donor integration showed complete loss of ATG5 protein expression (Supplementary Figure 2B, C). Functionally, these cells exhibited a complete loss of autophagy activity, as assessed by a standard autophagy flux assay (Supplementary Figure 2D). Re-introduction of an *ATG5* cDNA cassette into the safe-harbor locus *AAVS1* restored autophagy activity (Supplementary Figure 2E), confirming that the observed phenotype was due to specific loss of *ATG5*.

Next, we evaluated specificity through off-target analysis. To identify potential off-target cleavage events associated with our *ATG5* gRNA, we employed GUIDE-Seq, a technique that tags Cas9-induced double-strand breaks with deoxynucleotides, enabling their identification by next-generation sequencing (34). We mapped all double-strand breaks induced by plasmid electroporation in K562 cells, and detected GUIDE-Seq oligo tag integration at two off-target sites outside of the intended *ATG5* locus (Figure 1A; Supplementary Table 5). The predominant off-target site (“OT1”), which accounted for the majority of the off-target reads, was located in a non-coding RNA sequence on chromosome 4 (*LOC10537445*). A second, lower frequency site (“OT2”) was identified within the first intron of the *EBF2* gene on chromosome 8 (Supplementary Table 6). Next, we repeated the GUIDE-Seq process in CD3+ T cells from two healthy donors and observed consistent off-target editing at the same two sites, with OT1 being the dominant cleavage event (Figure 1B; Supplementary Table 6). To determine whether editing specificity could be improved, we tested a high-fidelity Cas9 variant (35) in the same healthy donor T cells. After performing HiFiCas9 RNP electroporation and amplicon sequencing, off-target editing at OT2 was eliminated (below the 0.1% limit of detection), while editing at OT1 was reduced to below 1% in both donors, which is comparable to previous off-target rates reported for clinical CAR-T products (36). Importantly, on-target editing efficiency at *ATG5* was maintained with HiFiCas9 (Figure 1C).

**Figure 1.**
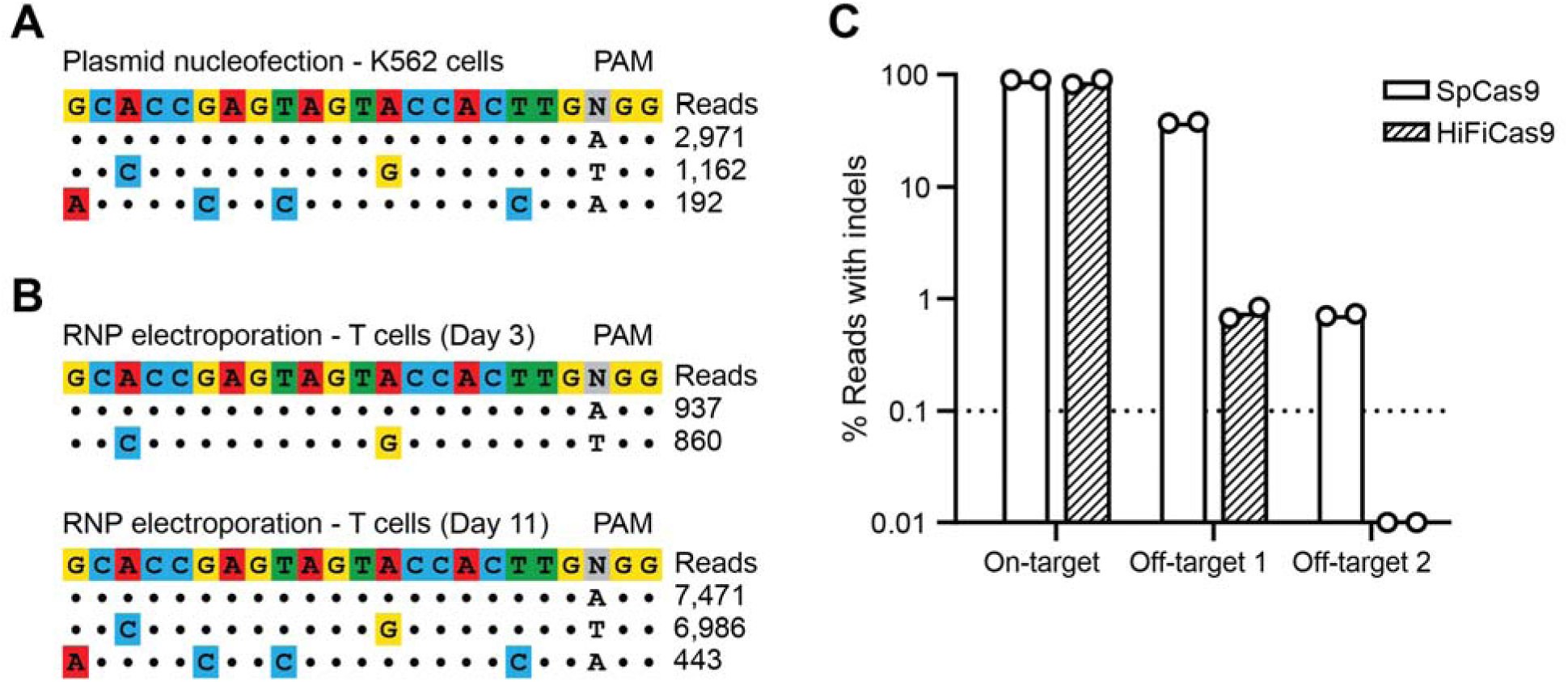
Editing at *ATG5* can be safely achieved with minimal off-target cleavage. **(A)** Allele plot showing off-target sites identified using GUIDE-Seq. K562 cells were electroporated with SpCas9-sgRNA vector targeting intron 2 of *ATG5* and GUIDE-Seq dsDNA, and genomic DNA was harvested three days post-electroporation. Dots represent matches with the target sequence, while mismatches are colored. **(B)** Same as (A) but for CD3+ T cells electroporated with SpCas9 RNPs targeting *ATG5* and GUIDE-Seq dsDNA. Genomic DNA was harvested on Day 3 (top) or Day 11 (bottom) post-electroporation. Day 3 read counts are from one of two different healthy donors. Day 11 read counts are from one healthy donor. **(C)** Indel quantification as determined by CRISPResso2 analysis from targeted amplicon sequencing of genomic DNA from T cells electroporated with SpCas9 or HiFiCas9 RNPs. The dotted line indicates a 0.1% limit of detection. Results are from n = 2 independent experiments performed with T cells from two different healthy donors.

After confirming feasibility and specificity of our editing approach, we designed AAV6 donors for CAR integration at *ATG5* and *AAVS1*. We used a published CAR targeting the antigen folate receptor alpha (αFR), which included a CD8 leader sequence, a single-chain variable fragment specific for human αFR, a CD8 hinge and transmembrane domain, a CD28 costimulatory domain, and a CD3ζ intracellular domain (37). Each donor included an elongation factor-1 alpha (EF1α) promoter upstream of the CAR expression cassette, along with a bovine growth hormone polyadenylation signal downstream of the CAR, all flanked by homology arms for either *ATG5* or *AAVS1* (Supplementary Figure 3A, B). *ATG5*-knockout and *AAVS1*-knockout αFR CAR-T cells were generated by RNP electroporation followed by incubation with concentrated AAV6 (3×10^5^ vg/cell) at a high cell density (4-5×10^6^ cells/ml). This method yielded greater than 70% CAR-positive cells (Figure 2A). Subsequent magnetic bead enrichment using an antibody specific for the CAR linker resulted in over 90% CAR-positive cells (Figure 2B). After ten days of *in vitro* expansion post-enrichment, CAR-T cells from both groups exhibited comparable viability and CD4:CD8 ratios (Figure 2C, D), although this protocol appeared to consistently favor expansion of CD4+ CAR-T cells (Figure 2D). Despite deletion of *ATG5*, there were no notable changes in CAR-T phenotype. Expression levels of CD137 and PD1 were similar between *ATG5*-and *AAVS1*-knockout cells (Figure 2E, F), as was the frequency of CCR7+/CD45RO+ central memory CAR-T cells (Figure 2G).

**Figure 2.**
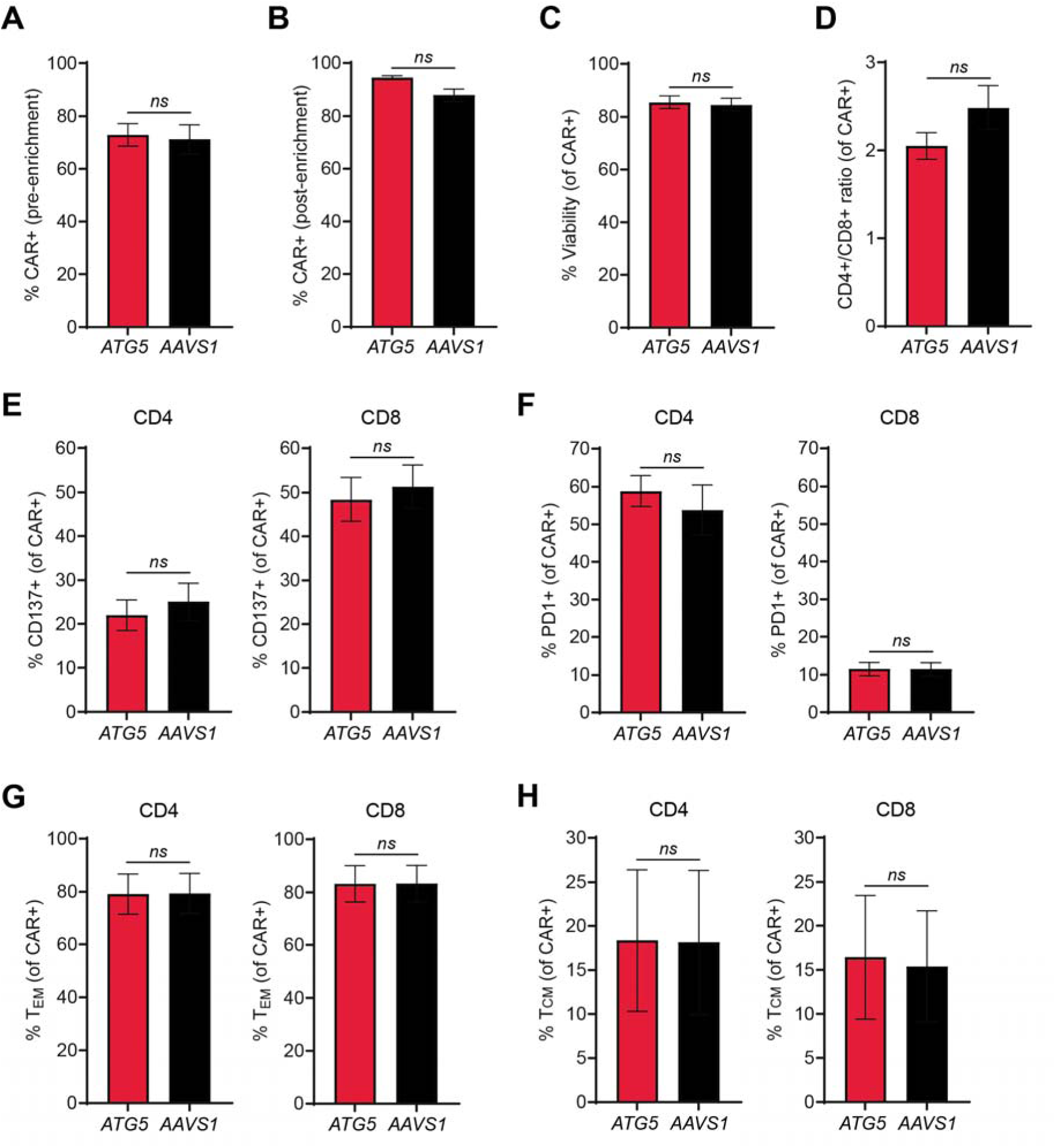
Integration at *ATG5* and *AAVS1* generates CAR-T cells with similar immune phenotypes. (A-B) Bar graphs showing percent CAR-positive cells pre-enrichment (A) and post-enrichment (B) for *ATG5*-knockout (red) and *AAVS1*-knockout (black) CAR-T cells. **(C-D)** Bar graphs showing percent viability (C) and ratio of CD4+ to CD8+ CAR-T cells (D). **(E-F)** Bar graphs showing percentage of CD4+ CAR-T cells (left) and CD8+ CAR-T cells (right) positive for (E) the activation marker CD137 and (F) the exhaustion marker PD1. **(G)** Bar graph showing percentage of CD45RO+/CCR7-effector memory CD4+ CAR-T cells (left) and CD8+ CAR-T cells (right). **(H)** Bar graph showing percentage of CD45RO+/CCR7+ central memory CD4+ CAR-T cells (left) and CD8+ CAR-T cells (right). Results are from two different healthy donors, with n = 2 independent experiments per donor. All error bars represent +/-SEM. Significance was determined by Welch’s t test: *ns*, not significant.

### Targeted CAR integration at *ATG5* metabolically reprograms CAR-T cells

In contrast to the similar immune phenotypes exhibited by *ATG5*-knockout and *AAVS1*-knockout CAR-T cells, targeted metabolite profiling revealed striking differences in baseline metabolic state (Figure 3A). *ATG5*-knockout CAR-T cells were significantly enriched in glucose and glutamine, two critical nutrients for T cell function and survival (38–42), suggesting increased nutrient acquisition (Supplementary Table 7). However, downstream intermediates of glycolysis and the tricarboxylic acid (TCA) cycle, including lactate, malate, fumarate, and succinate, were markedly reduced relative to *AAVS1*-knockout controls (Supplementary Table 7), indicating a potential uncoupling of nutrient uptake from pathway utilization. Additionally, *ATG5*-knockout CAR-T cells displayed increased enrichment of tyrosine, methionine, leucine, and phenylalanine (Supplementary Table 7), essential amino acids imported via the L-type amino acid transporter LAT1 (43). These findings align with recent evidence that *ATG5* regulates surface expression of nutrient transporters (44), and collectively support a model in which disruption of *ATG5* reprograms CAR-T cell metabolism by enhancing nutrient uptake rather than catabolism.

**Figure 3.**
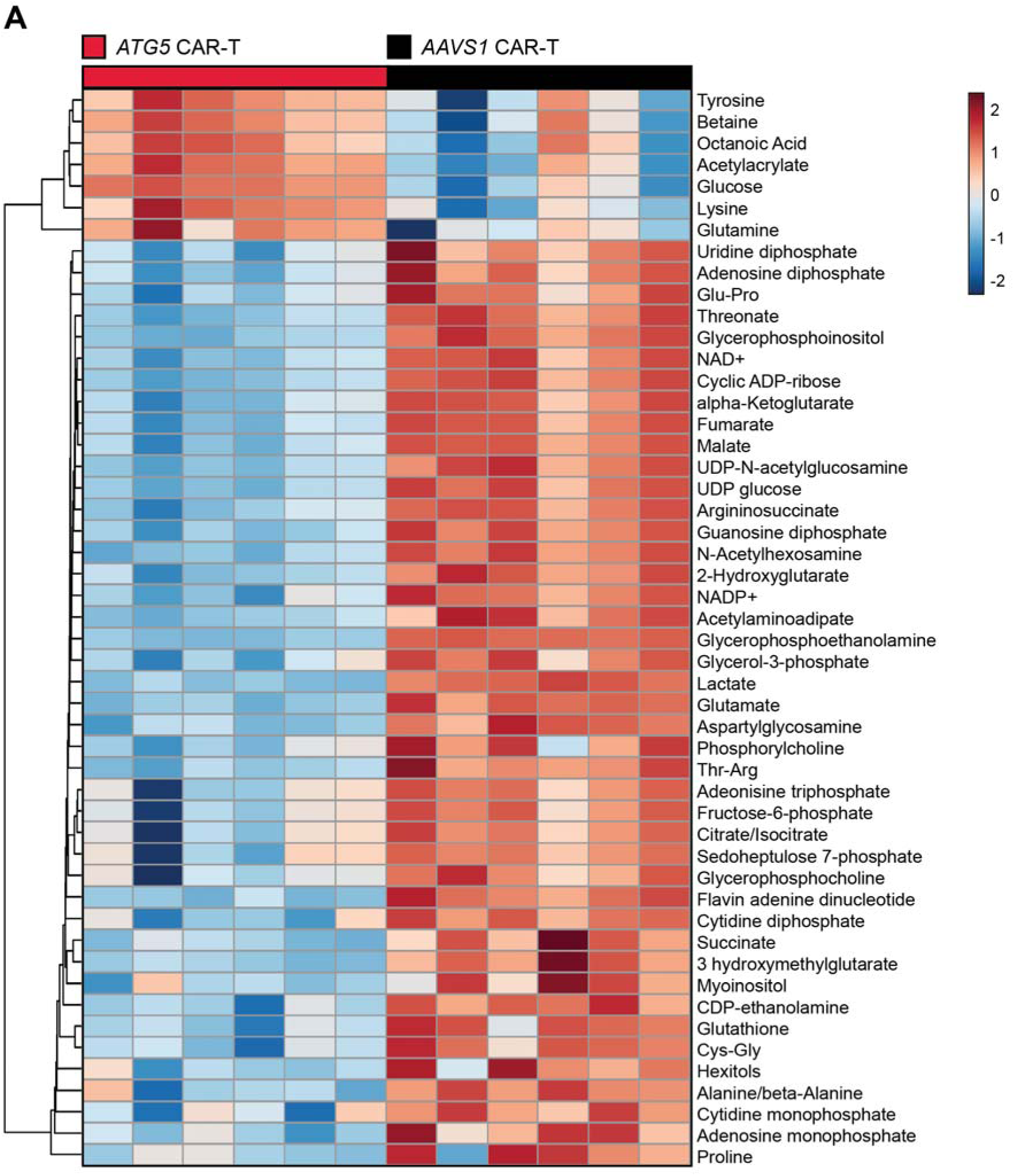
***ATG5*-knockout CAR-T cells shift to reduced catabolic metabolism. (A)** Heatmap of normalized metabolite abundances for the top 50 most significant metabolites, with dendrograms showing single-linkage clustering of Euclidean distances between metabolites. Metabolites were identified by liquid chromatography mass spectrometry analysis of enriched CAR-positive cells ten days post-enrichment. Results are from n = 2 healthy donors, 3 technical replicates per donor. Significance was determined by ANOVA (p < 0.05).

### *ATG5* integration enhances CAR-T effector function and cytotoxicity under immune-suppressive conditions

To determine whether the distinct metabolic profiles of *ATG5*-knockout CAR-T cells conferred functional advantages, we evaluated glucose uptake using an orthogonal, flow cytometry-based assay with fluorescently-labelled glucose. Consistent with our metabolomic findings, *ATG5*-knockout CAR-T cells exhibited higher glucose uptake compared to *AAVS1*-knockout controls (Figure 4A). Given previous evidence linking glucose uptake with effector function (38,45), we measured intracellular cytokine production and observed markedly elevated IFN-γ expression in both CD4+ and CD8+ *ATG5*-knockout CAR-T cells (Figure 4B). Enhanced cytokine production correlated with superior cytolytic activity across a range of effector-to-target cell ratios in standard *in vitro* killing assays against human αFR-expressing SKOV3 cells (Figure 4C).

**Figure 4.**
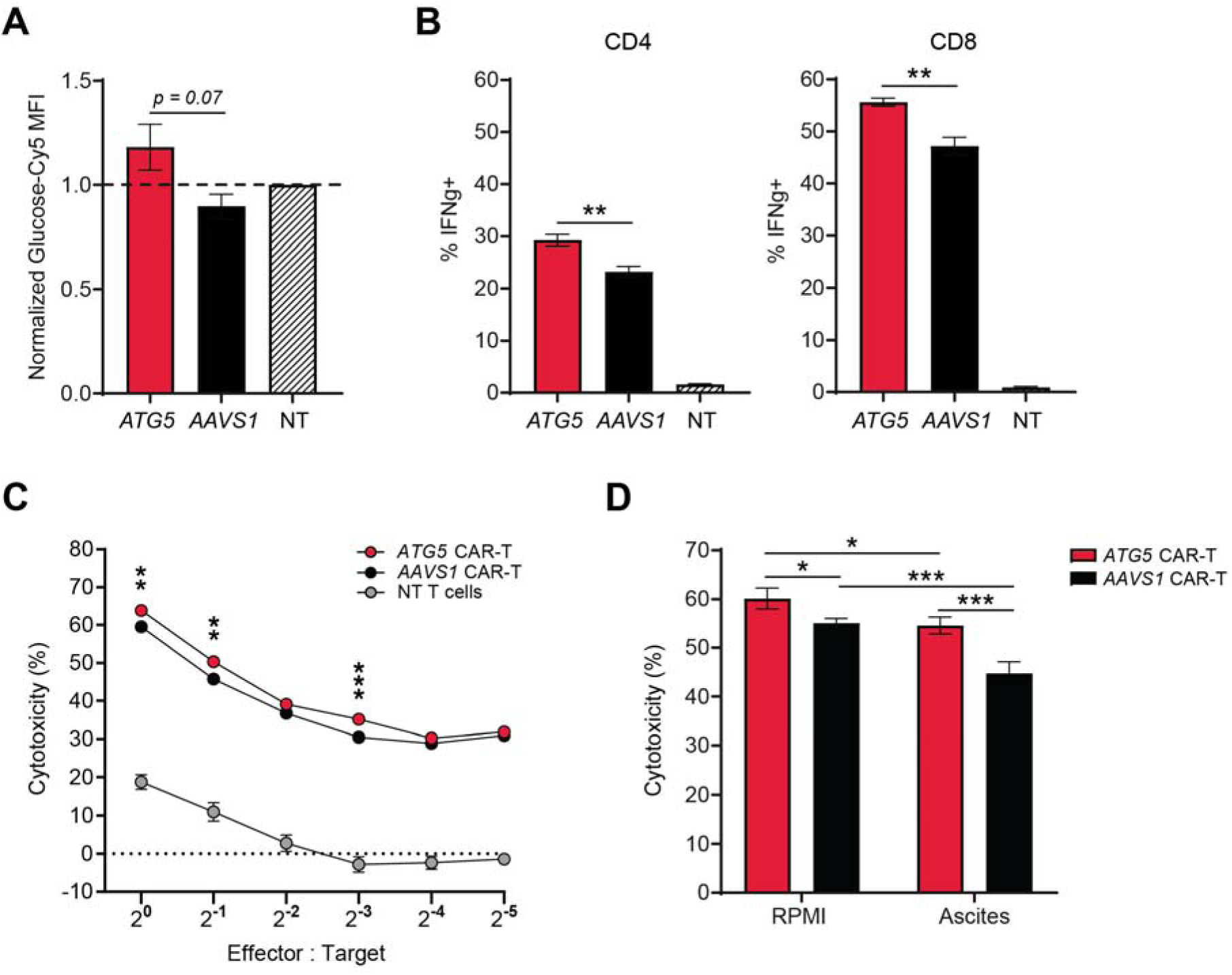
Integration at *ATG5* enhances glucose uptake and effector function of CAR-T cells. **(A)** Bar graph showing normalized glucose uptake for *ATG5*-knockout (red) and *AAVS1*-knockout (black) CAR-T cells. Glucose uptake was quantified as median fluorescence intensity (MFI) of Cy5-labelled glucose after a 30-minute incubation. MFI for each CAR-T group was normalized to MFI for non-transduced T cells within each individual experiment to allow for comparison between experiments. Results are from CAR-T cells made from two healthy donors, n = 2 independent experiments per donor, 2 technical replicates per experiment. **(B)** Bar graphs showing percentage of CD4+ CAR-T cells (left) or CD8+ CAR-T cells (right) positive for the cytokine interferon gamma after 24-hour co-culture with SKOV3 ovarian tumor cells at a 2:1 effector-to-target cell ratio. Non-transduced T cells were included as an assay control. Results are from one healthy donor, n = 2 independent experiments, 2 technical replicates per experiment. **(C)** Scatter plot showing percent cytotoxicity of luciferase-expressing SKOV3 ovarian tumor cells after 24-hour co-culture with CAR-T cells at the indicated E:T ratios. Assays were performed in RPMI. Results are from two healthy donors, n = 2 independent experiments per donor, 4 technical replicates per experiment. NT = non-transduced T cells. **(D)** Bar graphs showing percent cytotoxicity of luciferase-expressing SKOV3 cells after 24-hour co-culture with CAR-T cells at an E:T ratio of 1:1. Assays were performed in RPMI (left) or patient-derived ascites supernatant (right; n = 2 different ascites samples). Results are from CAR-T cells made from two healthy donors, n = 2 experiments per donor, 4 technical replicates per experiment. All error bars represent +/-SEM. P values were determined by Welch’s t test (A-C) or 2-way ANOVA (D): *p < 0.05, **p < 0.01, ***p < 0.001.

To test whether these functional enhancements would persist in a more physiologically relevant, immunosuppressive context, we repeated cytotoxicity assays in the presence of patient-derived ascites fluid, a pseudo *ex vivo* model of the ovarian tumor microenvironment. Ascites fluid contains numerous immune-inhibitory factors such as transforming growth factor beta (TGF-B) and interleukin-10 (IL-10), the suppressive metabolite kynurenine (22,46,47), and lower concentrations of glucose and glutamine as compared to standard cell culture medium (48).

As expected, overall CAR-T cytolytic activity was diminished in ascites fluid from two patients with HGSOC, consistent with a suppressed functional state. However, *ATG5*-knockout CAR-T cells remained robust, showing less than a 5% reduction in cytotoxicity, whereas *AAVS1*-knockout CAR-T cells experienced a significant functional decline (Figure 4D). These findings demonstrate that metabolic reprogramming via *ATG5* deletion enhances CAR-T cell effector function and confers resistance to TME-mediated immune suppression.

### *ATG5*-knockout CAR-T cells demonstrate superior tumor control *in vivo*

To determine whether the functional advantages of autophagy disruption observed in patient-derived ascites would translate to effective tumor control *in vivo*, we treated OVCAR3 tumor-bearing mice with 4×10^6^ *ATG5*-knockout or *AAVS1*-knockout CAR-T cells (Figure 5A). Both groups showed an early and immediate response following CAR-T infusion. However, the antitumor effect was markedly greater in mice treated with *ATG5*-knockout CAR-T cells, where the average reduction in tumor area over the first 17 days post-treatment was 3.693 mm^2^ per day (Figure 5B). In contrast, mice treated with *AAVS1*-knockout CAR-T cells experienced a much slower rate of tumor reduction within the same time period (1.058 mm^2^/day; Figure 5C).

**Figure 5.**
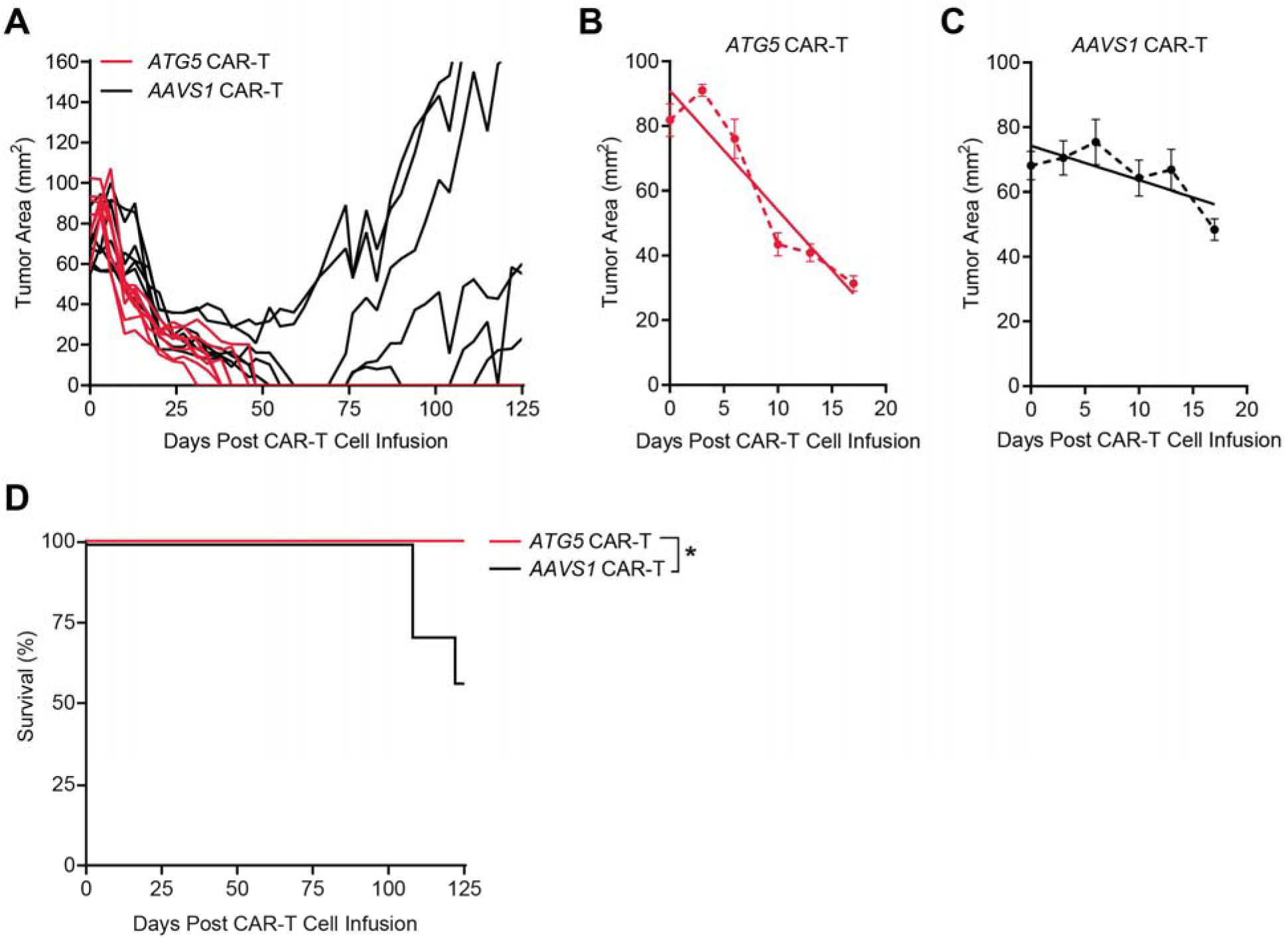
***ATG5*-knockout CAR-T cells exhibit immediate and long-lasting tumor control *in vivo*.** NSG mice were subcutaneously implanted with 5×10^6^ OVCAR3 tumor cells on the left flank. *ATG5*-integrated (n=8) or *AAVS1*-integrated (n=7) were intravenously administered 46 days after tumor implantation. Tumors were measured twice per week using digital calipers, and mice were euthanized upon reaching humane endpoint (tumor area greater than 160mm^2^) or at termination of study (125 days after CAR-T infusion). **(A)** Growth curve showing tumor area over time following treatment with 4×10^6^ *ATG5*-integrated (red) or *AAVS1*-integrated (black) CAR-T cells (one line = one mouse). **(B-C)** Scatter plots showing average tumor area for the first 17 days of the experiment shown in (A). Solid lines indicate the line of best fit for each group. Error bars represent +/-SEM. **(D)** Kaplan-Meier survival curve for the experiment shown in (A). Significance was determined by log-rank test: *p < 0.05.

Notable differences in therapeutic outcomes became more pronounced as the study progressed. By day 48, *ATG5*-knockout CAR-T cells had fully eliminated tumors in all treated mice, with no recurrence observed through study termination at day 125. By comparison, *AAVS1*-knockout CAR-T cells showed limited durability, with significantly reduced survival and recurrent tumors in the majority of remaining mice at study endpoint (Figure 5A, D). These findings demonstrate that the metabolic adaptations conferred by *ATG5* deletion are potent and sustained, thereby enabling CAR-T cells to achieve complete and durable tumor clearance in an ovarian cancer model.

## DISCUSSION

While the precise mechanisms limiting CAR-T function in ovarian cancer remain poorly defined, mounting evidence suggests that the TME imposes profound metabolic constraints that suppress effector T cell activity. We and others have previously shown that ovarian tumors are characterized by nutrient depletion, accumulation of suppressive metabolites, and widespread hypoxia, all of which impair cytokine production and cytolytic capacity (21,22,48). Therefore, the development of successful T cell-based therapies for ovarian cancer requires strategies to overcome downregulation of effector responses due to metabolic suppression in the TME. Here, we demonstrate that targeted disruption of the essential autophagy gene *ATG5* metabolically reprograms αFR CAR-T cells for improved function under immune-suppressive conditions, leading to sustained rejection of ovarian tumors *in vivo*.

CAR-T therapy has yet to deliver meaningful clinical benefit to patients with ovarian cancer (49). However, preclinical models consistently show that ovarian tumors are susceptible to CAR-T-mediated lysis (50,51), indicating that the barrier to efficacy may not lie in antigen escape, but in other features of the TME. Interestingly, while there is a well-recognized prognostic benefit associated with increased intratumoral T cells in HGSOC (52–54), recent evidence suggests that many of these infiltrating T cells are exhausted or dysfunctional (55–57). To that end, strengthening the early effector response, rather than focusing on long-term persistence, presents an alternative strategy for increasing CAR-T efficacy in ovarian cancer. Indeed, our *in vivo* study revealed notable differences in efficacy, where treatment with *ATG5*-knockout CAR-T cells elicited an immediate reduction in tumor growth, followed by rapid tumor clearance and complete remission in all mice. In contrast, *AAVS1*-knockout CAR-T cells were less effective in the first three weeks post-treatment and failed to completely control tumors, leading to multiple recurrences. Taken together, these findings suggest that there is a correlation between the magnitude of the early effector response and overall *in vivo* efficacy, and underscore the therapeutic potential of *ATG5* deletion as a strategy to improve CAR-T outcomes in ovarian cancer.

Metabolically, *ATG5*-knockout CAR-T cells are highly enriched in glucose, glutamine, and other essential amino acids as compared to control CAR-T cells. Glucose and amino acids are required for T cell activation, differentiation, and effector function (39,58,59), and depletion of these nutrients has well-documented deleterious effects on cytokine production and cytotoxicity (40,60,61). Therefore, the functional advantages conferred by *ATG5* deletion could be attributed to the development of a favorable metabolic state with increased capacity to support effector function. However, we did not observe corresponding increases in downstream intermediates of glucose or amino acid metabolism in *ATG5*-knockout CAR-T cells, which indicates that deletion of *ATG5* metabolically reprograms CAR-T cells through increased nutrient uptake, rather than catabolic flux. Given that several of the amino acids enriched in *ATG5*-knockout CAR-T cells are transported by the same amino acid transporter, LAT1 (43), a potential explanation for this pattern of metabolite enrichment is that deletion of *ATG5* alters the expression of nutrient transporters on the cell surface. This hypothesis is supported by recent studies showing that ATG5 interacts with the clathrin-mediated endocytic pathway to regulate vesicular trafficking of cell surface proteins (44), and with the ubiquitin-proteosome pathway to mediate degradation of c-Myc (62), a known positive regulator of LAT1 and other amino acid transporters (63).

Furthermore, a recent proteomic analysis demonstrated that autophagy preferentially degrades glucose and amino acid transporters, but this can be overcome by inhibiting vacuolar protein sorting 34 (VPS34), a key protein involved in autophagosome formation (30). These findings support a model in which deletion of *ATG5* enhances expression of key nutrient transporters, which in turn endows *ATG5*-knockout CAR-T cells with a functional advantage in nutrient-restricted conditions such as the ovarian TME.

However, while our study demonstrates that *ATG5*-knockout CAR-T cells exhibit increased effector cytokine production, the precise molecular mechanisms linking loss of autophagy with immuno-metabolic reprogramming remain undefined. Although several studies have shown that systemic loss of autophagy leads to changes in cytokine secretion by tumor-infiltrating T cells, the underlying pathways responsible for this effect are still unknown (64). Similarly, the direct downstream consequences of autophagy gene deletion on the regulation of amino acid availability require further delineation (65). Together, these collective findings highlight the need for further investigation into how autophagy orchestrates amino acid metabolism and effector programming in T cells.

A possible challenge to our approach is that loss of autophagy prevents *ATG5*-knockout CAR-T cells from forming canonical memory populations, which could impair long-term persistence in some contexts. However, we observed durable tumor control and long-term remission after a single infusion of *ATG5*-knockout CAR-T cells, which suggests that the absence of memory markers does not preclude sustained therapeutic benefit. Importantly, in the setting of solid tumors such as ovarian cancer, where CAR-T therapies have historically shown limited efficacy, the generation of highly potent effector cells capable of initiating robust anti-tumor responses is likely a critical determinant of therapeutic success. Thus, while the lack of memory may represent a trade-off, our data suggest that the enhanced effector function conferred by autophagy deletion may prove more beneficial for overcoming barriers to CAR-T activity in solid tumors.

## MATERIALS AND METHODS

### Study design

The aim of this study was to investigate the effects of autophagy deletion on CAR-T cell metabolism and function in the context of ovarian cancer immunotherapy. We used CRISPR-Cas9 to generate *ATG5*-knockout or *AAVS1*-knockout CAR-T cells made from two separate healthy donors, and quantified off-target editing with GUIDE-Seq. Using targeted metabolite profiling, spectral flow cytometry, and luminescence-based cytotoxicity assays, we showed that deletion of *ATG5* metabolically reprograms CAR-T cells to support enhanced effector function under immune-suppressive conditions. We demonstrated the relevance of these findings to ovarian cancer by conducting an *in vivo* study using immunodeficient mice implanted with OVCAR3 tumors, where *ATG5*-knockout CAR-T cells exhibited complete and lasting tumor control.

### Cell line culture

K562 human leukemia cells (CCL-243, ATCC) were grown in RPMI 1640 (Gibco) supplemented with 10% heat-inactivated fetal bovine serum (FBS; Sigma), 2 mM L-glutamine (ThermoFisher), and 1% penicillin-streptomycin (ThermoFisher). HEK293T cells (CRL-3216, ATCC) were grown in DMEM (Gibco) supplemented with 10% FBS and 1% penicillin-streptomycin. Luciferase-expressing SKOV3 human ovarian cancer cells (SKOV3-Luc, a gift from Dr. Julian Smazynski) were grown in McCoy’s 5A (Gibco) supplemented with 10% FBS, 1% penicillin-streptomycin, and 2 μg/ml puromycin (Gibco). OVCAR3 human ovarian cancer cells (HTB-161, ATCC) were grown in ATCC-formulated RPMI 1640 (Gibco) supplemented with 20% FBS, 0.01 mg/mL bovine insulin (Sigma), and 1% penicillin-streptomycin.

### T cell culture

CD3+ T cells were isolated from healthy donor peripheral blood mononuclear cells (PBMCs; STEMCELL) using positive selection CD3 microbeads (Miltenyi). T cells were activated with 25 μl Immunocult CD3/CD28 T cell activators (STEMCELL) per 1 million cells and cultured in Immunocult-XF medium (STEMCELL) supplemented with 1% penicillin-streptomycin, 2 mM L-glutamine, and 300 U/ml IL-2 (Peprotech).

### Collection and processing of ascites supernatant for cytotoxicity assays

Patient ascites was obtained through the BC Cancer Tumor Tissue Repository under University of British Columbia Biosafety (B23-0067) and REB (H18-01783) protocols. Ascites was centrifuged at 1500 rpm for 10 minutes at 4 °C, and the supernatant removed by pipetting and filtered through a 40 μm cell strainer before cryopreservation at-80 °C.

### Guide RNA screening

*ATG5*-targeting gRNA sequences were designed using the online tool CRISPOR (33) and cloned into SpCas9-sgRNA vector (derived from Addgene 48318). For gRNA screening, 200,000 K562 cells were electroporated with 500 ng or 750 ng SpCas9-sgRNA vector on the 4D-Nucleofector X Unit (Lonza) using the SF nucleofection kit and pulse code FF-120. Genomic DNA (gDNA) was extracted from cells three days post-electroporation using QuickExtract (Lucigen) as per the manufacturer’s protocol. The percentage of edited alleles was quantified using the Surveyor mutation detection kit (Transgenomics) as previously described (66). Samples were run on 10% PAGE gels, imaged with a ChemicDoc MP system (Bio-Rad), and images quantified using Image Lab software (Bio-Rad). After screening, the gRNA sequence ATG5-I2A-824 was selected for all further experiments. The gRNA targeting intron 1 of *AAVS1* was previously described in the literature (67). Both gRNA sequences can be found in Supplementary Table 1.

### Design and synthesis of genome editing components

*ATG5* and *AAVS1* gRNAs were synthesized by IDT, resuspended in nuclease-free water to 100 μM, and stored at-80 °C. CAR donor sequences (ATG5-EF1α-αFR, and AAVS1-EF1α-αFR) were designed around a published CAR construct (37). Each donor included an EF1α promoter upstream of the CAR, and a bovine growth hormone polyadenylation sequence downstream of the CAR. Full donor sequences were synthesized as gBlocks (IDT) and cloned into adeno-associated virus serotype 6 (AAV6) vectors via restriction cloning. Recombinant CAR-AAV6 was produced by the viral vector core at the Canadian Neurophotonics Platform, and the virus was resuspended in PBS 320 mM NaCl + 5% D-sorbitol + 0.001% pluronic acid and stored at - 80 °C until use. Guide RNA sequences can be found in Supplementary Table 1. Full donor sequences can be found in Supplementary Data File S1.

### T cell electroporation

To prepare RNPs, gRNAs (100 μM, IDT) were mixed in a 2:1:1 molar ratio with poly-l-glutamic acid (PGA; 100 mg/ml, Sigma) and SpCas9 or HiFiCas9 nuclease as indicated (10 mg/ml, both IDT), and incubated for 15 minutes at 37 °C. T cells were resuspended in electroporation buffer P3 (Lonza), mixed with RNPs at a ratio of 1×10^6^ cells/50 pmol RNP and electroporated in 16-well nucleocuvette strips on the 4D-Nucleofector X Unit (Lonza) using pulse code EO115. For RNP-only conditions, 80 μl of pre-warmed T cell medium (no IL-2) was added to each well immediately post-electroporation, and the nucleocuvette strip was returned to the incubator for 15 minutes at 37 °C. After 15 minutes, T cells were transferred to a 48-well plate at a density of 1×10^6^ cells/ml in complete T cell medium.

### Generation of *ATG5*-knockout and *ATG5*-complemented K562 clones

To generate *ATG5*-knockout K562 cell lines, wild-type K562 cells were electroporated with 350 ng SpCas9-sgRNA vector (derived from Addgene 48318) and 700 ng *ATG5*-mScarlet-I_NLS donor. Three days post-transfection, cells were single-cell sorted on mScarlet-I expression using a FACSAria cytometer (BD Biosciences) and expanded for 10 days in RPMI (Gibco). Single-cell clones were screened for tri-allelic integration using out-out PCR, and loss of ATG5 was verified by Western blot. To generate *ATG5*-complemented K562 cell lines, *ATG5*-knockout clone 5 was electroporated with 350 ng SpCas9-sgRNA vector and 700 ng *AAVS1*_Puro_PGK1_ATG5-cDNA donor (derived from Addgene 68375; (68)). Single-cell clones were picked and expanded for 10 days in methylcellulose-based semi-solid RPMI (Gibco) supplemented with 0.5 μg/ml puromycin.

### Autophagy flux assay

Wild-type, *ATG5*-knockout, and *ATG5*-complemented K562 cells were assessed for autophagy function using a standard flux assay. Cells were cultured for 20 hours at 37 °C in medium supplemented with 30 μM hydroxychloroquine and/or 200 nM rapamycin (both Cayman Chemicals). After culture, cells were pelleted and resuspended in RIPA buffer (50 mM Tris-HCl pH 7.4, 1% NP-40, 0.25% Na-deoxycholate, 150 mM NaCl, 1 mM EDTA) supplemented 1:100 with protease and phosphatase inhibitor cocktail (ThermoFisher). Cells were lysed for 30 minutes on ice, then centrifuged for 10 minutes at 13000 rpm and 4 °C. Lysates were mixed 1:10 with sample reducing agent and 1:4 with LDS buffer (both Invitrogen) and heated for 10 minutes at 70 °C before loading onto 4-12% gradient Bis-Tris gels (Invitrogen). Protein was transferred onto nitrocellulose membranes, blocked for 60 minutes in blocking buffer (LI-COR Biosciences), diluted 1:1 with PBS, and stained overnight at 4 °C with anti-ATG5, anti-LC3, anti-Tubulin, or anti-GAPDH as indicated. After washing, membranes were incubated for 1 hour at room temperature with anti-mouse or anti-rabbit secondary antibodies. All staining was performed in blocking buffer diluted 1:1 with PBS. A detailed list of all antibodies can be found in the Supplemental Material.

### GUIDE-Seq

GUIDE-Seq was performed as previously described (34,69). GUIDE-Seq oligos with 5’ phosphorylation and two phosphorothioate linkages between the last two nucleotides at the 3’ end were synthesized by IDT. For K562 cells, 2×10^5^ cells were electroporated with 750 ng SpCas9-sgRNA vector and 100 pmol of annealed GUIDE-Seq oligos. For T cells, 1×10^6^ cells were electroporated with 50 pmol SpCas9 RNP and 100 pmol annealed GUIDE-Seq oligos.

Genomic DNA was harvested using the QIAmp UCP DNA Micro Kit (QIAGEN) and sequenced by Sanger sequencing, and GUIDE-Seq oligo tag integration was confirmed using TIDE (70) and DECODR (71). The Next Generation Sequencing (NGS) library preparation procedure was modified from the original GUIDE-Seq procedure (72). In brief, end-repair and A-tailing were directly performed using the NEBNext UltraII DNA kit (New England Biolabs). A new custom universal Y-adapter based on Illumina TruSeq sequences was designed to include a Unique Molecular Index (UMI) in the sequencing read instead of the index read. This adapter was ligated as described in the NEBNext Ultra II procedure using a final concentration of 133 nM. Ligated DNA was purified using AMPure XP beads (Beckman-Coulter) at a 0.9X volumetric ratio, then PCR-amplified with either Minus or Plus primers using Q5 DNA polymerase (New England Biolabs). PCR products from reaction 1 were purified with 0.9X AMPure XP beads and quantified by Qubit dsDNA HS assay (Invitrogen). A second PCR amplification was performed using 10 ng of purified DNA from PCR1 and Illumina TruSeq dual-indexing primers. PCR products from reaction 2 were purified using 0.85X AMPure XP beads, quantified by Qubit dsDNA HS assay, and run on a Bioanalyzer High Sensitivity DNA chip (Agilent). Samples were pooled in equimolar amounts and sequenced on a 1.4% (35M read block) of a NovaSeq S4 PE150 lane (Illumina) at the Centre d’Expertise et de Services Génome Québec (Montréal, QC). Sequencing data from the GUIDE-Seq procedure was analyzed as described in the Supplemental Material. Detailed GUIDE-Seq results and sequences of all primer and adapter sequences can also be found in the Supplemental Material.

### CAR-T cell production

T cells were electroporated with *ATG5* or *AAVS1* RNPs as described in the previous section. Immediately following electroporation, T cells were transferred from the nucleocuvette strip to a 96-well round-bottom plate containing CAR-AAV6 diluted in T cell medium at an MOI of 3 x10^5^ vg/cell. The plate was returned to the incubator for 6-8 hours incubation with AAV6 at a high cell density, after which T cells were transferred to a 48-well plate at a density of 1×10^6^ cells/ml in complete T cell medium. Five days post-electroporation, edited cells were collected and stained for 30 minutes at room temperature with AF647-conjugated anti-G4S linker antibody (Cell Signaling) diluted 1:50 in PBS. After staining, cells were incubated for 15 minutes at 4 °C with anti-AF647 microbeads (Miltenyi) in cell-enrichment buffer (PBS supplemented with 0.5% heat-inactivated human serum (Sigma)), then passed through an MS column (Miltenyi) for enrichment. CAR-positive cells were plated at 500,000 to 800,000 cells/ml in fresh medium containing 20 μL/ml Immunocult CD3/CD28/CD2 T cell activators (STEMCELL). T cell activators were diluted by addition of fresh medium after 2 days of re-stimulation. For the remaining culture period, CAR-T cells were maintained at 500,000 to 1 million cells/ml and transferred to larger culture vessels as needed.

### LC-MS metabolite profiling

To prepare CAR-T cells for metabolite profiling, triplicate samples of 2 million cells were collected from each group, washed once with ice-cold saline solution, and vigorously resuspended in 500 μl of 80% methanol before snap freezing in liquid nitrogen. Samples were subjected to 3 freeze-thaw cycles, then centrifuged at 14,000 rpm for 15 minutes at 4 °C. Metabolite-containing supernatants were evaporated until dry, reconstituted in 100 μl of acetonitrile/water 80:20, vortex-mixed, and centrifuged to remove debris. Metabolites were separated by LC-MS and analyzed on the Thermo Scientific Orbitrap Exploris 480 high-resolution mass spectrometer in full scan mode as previously described (73–75). Chromatogram review and peak area integration were performed using TraceFinder software version 5.1 (Thermo Scientific). Peak areas were normalized to total ion count (TIC) to account for variations in sample handling and instrument performance. After TIC-normalization, multiple experiments were merged into the same dataset and corrected for batch effects using EigenMS (76). The normalized data were filtered by interquartile range, log-transformed, and auto-scaled using MetaboAnalyst (77). To visualize differences in metabolite abundance across CAR-T groups, heatmap dendrograms were constructed using a single-linkage method to cluster Euclidean distances among metabolites. To determine the role of autophagy deletion, linear modeling was performed using the Statistical Analysis [metadata table] module in MetaboAnalyst, with autophagy status as the primary variable and donor as a covariate for adjustment. The output of this analysis can be found in the Supplemental Material.

### Flow cytometry analysis

For glucose uptake analysis, 1×10^6^ CAR-T cells per condition were stained for 30 minutes at 37 °C in TexMACS (Miltenyi) containing 0.1 μM Glucose-Cy5 (Sigma). Cells were washed, stained with fixable viability dye eF506 for 15 minutes at 4 °C, washed again, then stained for 20 minutes at room temperature with CD3-BV750 diluted in PBS. For immune phenotyping, CAR-T cells were stained with viability-eF506 for 15 minutes at 4 °C, then washed and resuspended in Human TruStain FcX blocking solution (Biolegend) and Brilliant Stain Buffer Plus (BD Biosciences) for 10 minutes at room temperature. After blocking, cells were stained for 20 minutes at room temperature with the following antibody cocktail diluted in PBS: CD3-BV750 (Biolegend), CD4-AF700 (Biolegend), CD8-PerCP (Biolegend), CAR-AF647, CD45RO-PE-Cy7 (Invitrogen), CCR7-APC/Fire750 (Biolegend), CD137-BV605 (Biolegend), and CD279-BV421 (Biolegend). For intracellular staining, CAR-T cells were co-cultured for 24 hours with SKOV3 human ovarian cancer cells at an effector-to-target cell ratio of 2:1. In the last 4 hours of co-culture, GolgiStop (BD Biosciences) was added to each well as per the manufacturer’s recommended dilution. At the end of the co-culture period, cells were collected and stained with viability-eF506 for 15 minutes at 4 °C, then washed and stained for 20 minutes at room temperature with the following antibody cocktail diluted in PBS: CD3-BV750, CD4-AF700, CD8-PerCP, and CAR-AF647. After surface staining, cells were resuspended in Cytofix/Cytoperm Solution (BD Biosciences) for 20 minutes at 4 °C, washed twice with Perm/Wash Buffer (BD Biosciences), and then stained for 30 minutes at 4 °C with IFN-γ-BV421 (Biolegend) diluted in Perm/Wash Buffer. All cells were resuspended in PBS prior to acquisition on a Cytek Aurora spectral flow cytometer. Cytometry data were analyzed using SpectroFlo (Cytek) and FlowJo v10.10 (BD Life Sciences). A detailed list of all antibodies can be found in the Supplemental Material.

### Cytotoxicity assays

Cytotoxicity assays were conducted in assay medium (RPMI 1640 supplemented with 5% human serum (Sigma), 12.5 mM HEPES (Gibco), 2 mM L-glutamine, and 1% penicillin-streptomycin) or patient-derived ascites supernatant. CAR-T cells were co-cultured in a 96-well plate with luciferase-expressing SKOV3 human ovarian cancer cells (SKOV3-Luc) at an effector-to-target ratio of 1:1. SKOV3-Luc cells were cultured alone to determine the maximum luciferase expression (RLU_max_), while medium-or ascites-only wells were used to account for background luminescence (RLU_background_). After 24 hours of co-culture, D-Luciferin (1/20; Revvity Health Sciences) was added to each well, mixed by pipetting, and the plate was incubated in the dark for 5 minutes. Luminescence was measured on a Varioskan Lux plate reader (Thermo Scientific), and specific tumor cell lysis calculated as (RLU_sample_/((RLU_max_-RLU_background_)) x 100.

### Animal study

The animal study was approved by the University of Victoria’s Animal Care Committee (AUP 2022-11) and performed in accordance with the Canadian Council for Animal Care guidelines. 6-8 weeks old female NOD/SCID/IL-2Rγ-null (stock #005557) mice were acquired from Jackson Laboratory. After a two-week acclimation period, each mouse was subcutaneously injected on the left flank with 5×10^6^ OVCAR3 tumor cells. Once tumors reached an average area of 70-80 mm^2^, mice were treated with 4×10^6^ CAR-positive cells (n=7 or 8 mice per CAR-T group) or 4×10^6^ non-transduced T cells (n=5 mice), administered in 100 μl PBS via tail vein injection. Mice were randomly assigned to each group to ensure similar distribution of tumor size. Tumors were measured twice per week using digital calipers, and area (length x width) was used to quantify tumor burden. Tumor measurements were conducted by technical staff who were blinded to the treatment intervention. Mice were euthanized upon reaching humane endpoint (tumor area greater than 160 mm^2^) or at termination of study, and overall survival was assessed by Kaplan-Meier analysis. All CAR-T cells for this study were manufactured using HiFiCas9.

### Quantification and statistical analysis

Statistical details for each experiment are indicated in the associated figure legends. Statistical analysis of LC-MS data was performed using MetaboAnalyst (77). All other data analyses were performed using GraphPad Prism version 10.1.1. Data are presented as +/-SEM, and error bars represent the SEM of two biological replicates, unless otherwise noted in the figure legend.

Significance was determined by Welch’s *t* test, one-way ANOVA, two-way ANOVA, linear regression followed by analysis of covariance (ANCOVA), or log-rank test: * *p* <0.05; ** *p* < 0.01; *** *p* < 0.001; **** *p* < 0.0001; *ns*, not significant.

## Supplementary Materials

Materials and Methods

Tables S1 to S8

Figures S1 to S3

Data files S1 and S2

## DECLARATIONS

## Acknowledgements

The authors would like to thank the Viral Vector Core at Neurophotonics for producing AAV6 for all genome-editing experiments, as well as Ahmed Olodo, Alison Bohnet, Celestine Aniugwu, Jennifer MacDonald, Laura Giguere, and the Animal Care staff at the University of Victoria for providing technical support for the animal study. The authors gratefully acknowledge the animals used in this study and recognize their invaluable contribution to advancing scientific knowledge.

## Author contributions

Conceptualization: GAC, SL, YD, JJL

Methodology: GAC, SL, RJD, PHW, YD, JJL

Investigation: GAC, SL, LGZ, JSP, TS

Funding acquisition: YD, JJL

Writing – original draft: GAC, JJL

Writing – review and editing: all

## Ethics approval

GAC and JJL have Research Ethics Board (REB) approval through the University of Victoria (18–1275) and the University of British Columbia (H18-01783), and Animal Ethics approval through the University of Victoria (2022-11). PHW holds REB approval from the BC Cancer Research Ethics Board for the patient biospecimen and data collections undertaken by the BC Cancer Tumor Tissue Repository (H18-003344 and H07-00463). The BC Cancer Tumor Tissue Repository is certified through the Canadian Tissue Repository Network certification program (https://biobanking.org/locator).

## Availability of data and material

All data needed to evaluate the conclusions in the paper are present in the paper or in the Supplementary Materials (data files S1 and S2).

