## Supplemental Material for "Autophagy disruption primes CAR-T cell metabolism for sustained rejection of ovarian tumors"

**SUPPLEMENTARY MATERIAL**

**Materials and Methods**

**Supplementary Table 1.** Guide RNA sequences

**Supplementary Table 2.** Antibodies for Western blot

**Supplementary Table 3.** PCR primer sequences

**Supplementary Table 4.** GUIDE-Seq adapter and primer sequences

**Supplementary Table 5.** GUIDE-Seq genomic targets

**Supplementary Table 6.** GUIDE-Seq results

**Supplementary Table 7.** Statistical analysis of metabolite profiling data

**Supplementary Table 8.** Antibodies for flow cytometry

**Supplementary Figure 1.** Guide screening identifies a highly active gRNA at intron 2 of *ATG5*

**Supplementary Figure 2.** Editing at *ATG5* eliminates functional autophagy

**Supplementary Figure 3.** Generating *ATG5*-knockout and *AAVS1*-knockout CAR-T cells

**Materials and Methods**

**Analysis of GUIDE-Seq sequencing data**

Sequencing data from the GUIDE-Seq procedure was analyzed using the guideseq_ibis pipeline (https://github.com/enormandeau/guideseq_ibis). Raw reads were trimmed for quality using trimmomatic (v0.36, min_length 100, crop_length 200), and reads containing the expected alien sequence (maximum hamming distance of 1) and only one copy of the GUIDE-Seq ODN sequence (maximum hamming distance of 6) were retained. For these reads, the UMIs and first eight nucleotides were used to rename the sequence, and the reads were mapped onto the latest GRCh38 human genome assembly with bwa (v0.7.17-r1188, -T 10) and samtools (v1.12, -S -q 1 -F 4 -F 256 -F 2048). The alignment sam files were then sorted by chromosome name and position. To identify double stranded breaks (DSBs), alignment sites were scanned for peaks of coverage in decreasing order of depth of coverage, and pairs of ODN+ and ODN- were reported until the pairs did not meet the minimum coverage threshold (min_length 100, min_coverage 50, window_size 10, position_error 5, bin_size 10000). To account for small errors in sequencing and mapping, read counts were collected within “window_size” nucleotides around each peak, and pairs of ODN sequences were only kept if they fell within “position_error” nucleotides of their relative expected positions. Identified sequences on-target and off-target were reported for each sample with their chromosome and position localisation, the identifier and name of any gene they overlapped, and metrics about the counts of ODN+ and ODN- reads.

**Supplementary Table 1.** Guide RNA sequences.

| **Sequence ID** | **DNA/RNA Sequence** |
| --- | --- |
| ATG5-I2A-824 | GTAATCTGACTTTCCGTAGC |
| AAVS1-I1 | GGGGCCACTAGGGACAGGAT |

**Supplementary Table 2.** Antibodies for Western blot.

| **Antibody** | **Dilution** | **Company** | **Cat. Number** |
| --- | --- | --- | --- |
| ATG5 | 1/500 | Sigma | A0731 |
| LC3 | 1/500 | Novus | NB100-2220 |
| Tubulin | 1/1000 | Santa Cruz | SC-32293 |
| GAPDH | 1/1000 | Novus | NB300-221 |
| Mouse IgG | 1/10000 | Invitrogen | A21240 |
| Rabbit IgG | 1/5000 | Invitrogen | A21109 |

**Supplementary Table 3.** PCR primer sequences.

| **Primer ID** | **Sequence** | **Product size (bp)** |
| --- | --- | --- |
| ATG5-Out-F  ATG5-Out-R | CATCAGCTATGGTGCCTTCTTG  CAACTTACGCTCTAGTGCTCAC | 695 bp (WT)  or 758 bp (KI) |
| ATG5-I2A-824-TIDE-F  ATG5-I2A-824-TIDE-R | GACTTGCAGGTGTGAGTTAATGG  GAGTACCAGTAACAAGATTGCCG | 624 bp |
| AAVS1-I1-TIDE-F  AAVS1-I1-TIDE-R | TTCGGGTCACCTCTCACTCC  GGCTCCATCGTAAGCAAACC | 469 bp |
| ATG5-I2A-824-OT1-F  ATG5-I2A-824-OT1-R | AAAATCCTTCCCCGTTCCTCGAG  CACATTAATAGTGCACGTGACCCTC | 773 bp |
| ATG5-I2A-NGS-F | ACACTCTTTCCCTACACGACGCTCTTCCGATCTTGAAGTCTGCCCTTTGCTTTCC | 392 bp with extension  461 bp after indexing |
| ATG5-I2A-NGS-R | GTGACTGGAGTTCAGACGTGTGCTCTTCCGATCTGAGTACCAGTAACAAGATTGCCG |  |
| ATG5-824-OT1-NGS-F | ACACTCTTTCCCTACACGACGCTCTTCCGATCTTTTCAAGGCAAGTGACCGGAATG | 437 bp with extension  506 bp after indexing |
| ATG5-824-OT1-NGS-R | GTGACTGGAGTTCAGACGTGTGCTCTTCCGATCTCACATTAATAGTGCACGTGACCCTC |  |
| ATG5-824-OT2-NGS-F | ACACTCTTTCCCTACACGACGCTCTTCCGATCTGGGTTCTTACCCACTACCTTTC | 366 bp with extension  435 bp after indexing |
| ATG5-824-OT2-NGS-R | GTGACTGGAGTTCAGACGTGTGCTCTTCCGATCTCACTCACCATCCCGTCTTTAG |  |

**Supplementary Table 4.** GUIDE-Seq adapter and primer sequences.

Phosphorothioate linkages are illustrated with a star (*).

| **Primer or adapter ID** | **Sequence** |
| --- | --- |
| dsODN_Fwd | /5Phos/GTTTAATTGAGTTGTCATATGTTAATAACGGT*A*T |
| dsODN_Rev | /5Phos/ATACCGTTATTAACATATGACAACTCAATTAA*A*C |
| GUIDE-Seq_Adapter_Top | ACACTCTTTCCCTACACGACGCTCTTCCGATCTNNWNNWNNCCATCTCATCCCTGC*T |
| GUIDE-Seq_Adapter_Bot | /5Phos/GCAGGGATGAGAT*G*G |
| GUIDE-Seq_PCR1_Common | ACACTCTTTCCCTACACGACGCTCTTCCGATCT |
| GUIDE-Seq_PCR1_Plus | GTGACTGGAGTTCAGACGTGTGCTCTTCCGATCTCGTTATTAACATATGACAACTCAATTAAAC |
| GUIDE-Seq_PCR1_Minus | GTGACTGGAGTTCAGACGTGTGCTCTTCCGATCTTTGAGTTGTCATATGTTAATAACGGTA |

**Supplementary Table 5.** GUIDE-Seq genomic targets.

Nucleotide mismatches are highlighted in red.

| **Guide** | **Genomic target** | **Sequence** | **PAM** | **Mismatches** |
| --- | --- | --- | --- | --- |
| On-target | *ATG5* intron 2 | GCACCGAGTAGTACCACTTG | AGG | - |
| Off-target 1 | *LOC105374445* ncRNA | GC**C**CCGAGTAGT**G**CCACTTG | TGG | 2 |
| Off-target 2 | *EBF2* intron 1 | **A**CACC**C**AG**C**AGTACCAC**C**TG | AGG | 4 |

**Supplementary Table 6.** GUIDE-Seq results.

| **K562 cells** | | | | | | | |
| --- | --- | --- | --- | --- | --- | --- | --- |
| **Target ID** | **Chromosome/Gene** | | **Site 1** | **Site 2** | **Count 1** | **Count 2** | **Total reads** |
| On-target | Chr6 – *ATG5* | | 106314933 | 106315033 | 1020 | 1951 | 2971 |
| Off-target 1 | Chr4 – N/A | | 48270100 | 48270200 | 373 | 789 | 1162 |
| Off-target 2 | Chr8 – *EBF2* | | 26042535 | 26042635 | 97 | 95 | 192 |
| **T cells – Donor 1 (Day 3)** | | | | | | | |
| **Target ID** | | **Chromosome/Gene** | **Site 1** | **Site 2** | **Count 1** | **Count 2** | **Total reads** |
| On-target | Chr6 – *ATG5* | | 106314933 | 106315033 | 525 | 784 | 1309 |
| Off-target 1 | Chr4 – N/A | | 48270100 | 48270200 | 291 | 851 | 1142 |
| Off-target 2 | Chr8 – *EBF2* | | 26042535 | 26042635 | - | - | - |
| **T cells – Donor 2 (Day 3)** | | | | | | | |
| **Target ID** | | **Chromosome/Gene** | **Site 1** | **Site 2** | **Count 1** | **Count 2** | **Total reads** |
| On-target | | Chr6 – *ATG5* | 106314933 | 106315033 | 343 | 594 | 937 |
| Off-target 1 | | Chr4 – N/A | 48270100 | 48270200 | 195 | 665 | 860 |
| Off-target 2 | | Chr8 – *EBF2* | 26042535 | 26042635 | **-** | **-** | **-** |
| **T cells – Donor 2 (Day 11)** | | | | | | | |
| **Target ID** | | **Chromosome/Gene** | **Site 1** | **Site 2** | **Count 1** | **Count 2** | **Total reads** |
| On-target | Chr6 – *ATG5* | | 106314933 | 106315033 | 2966 | 4505 | 7471 |
| Off-target 1 | Chr4 – N/A | | 48270100 | 48270200 | 1940 | 5046 | 6986 |
| Off-target 2 | Chr8 – *EBF2* | | 26042535 | 26042635 | 188 | 255 | 443 |

**Supplementary Table 7.** Statistical analysis of metabolite profiling data.

Analysis was performed on normalized metabolite abundances obtained in technical triplicate from n = 2 independent healthy donors (FC = knockout – wildtype). Significance was reported as P_adj_<0.05.

| **Metabolite** | **logFC** | **AveExpr** | **t** | **P.Value** | **adj.P.Val** |
| --- | --- | --- | --- | --- | --- |
| Glycerophosphoethanolamine | -1.9063 | 1.50E-15 | -9.7328 | 2.66E-07 | 2.18E-05 |
| Lactate | -1.8927 | 1.04E-15 | -9.2071 | 5.01E-07 | 2.18E-05 |
| Glutamate | -1.873 | -3.24E-15 | -8.5568 | 1.14E-06 | 3.30E-05 |
| Glycerophosphoinositol | -1.8521 | 2.66E-15 | -7.9719 | 2.47E-06 | 5.37E-05 |
| N-Acetylhexosamine phosphate | -1.8444 | 9.25E-18 | -7.7662 | 3.28E-06 | 5.57E-05 |
| Flavin adenine dinucleotide | -1.8392 | -1.39E-17 | -7.6515 | 3.84E-06 | 5.57E-05 |
| Cyclic ADP-ribose | -1.8298 | 1.11E-15 | -7.4345 | 5.22E-06 | 5.93E-05 |
| UDP-glucose | -1.8233 | 2.43E-15 | -7.324 | 6.12E-06 | 5.93E-05 |
| UDP-N-acetylglucosamine | -1.8237 | 5.87E-16 | -7.3073 | 6.27E-06 | 5.93E-05 |
| Guanosine diphosphate mannose | -1.82 | 1.85E-17 | -7.2495 | 6.82E-06 | 5.93E-05 |
| Threonate | -1.8171 | 9.11E-16 | -7.1816 | 7.52E-06 | 5.95E-05 |
| Aspartylglycosamine | -1.8117 | -9.25E-18 | -7.0759 | 8.78E-06 | 6.37E-05 |
| 2-Hydroxyglutarate | -1.7981 | 9.25E-18 | -6.8207 | 1.28E-05 | 8.59E-05 |
| Malate | -1.7926 | 3.70E-16 | -6.7257 | 1.48E-05 | 9.21E-05 |
| Argininosuccinate | -1.7764 | 4.63E-18 | -6.5901 | 1.82E-05 | 0.000101 |
| NAD+ | -1.7838 | -8.19E-16 | -6.581 | 1.85E-05 | 0.000101 |
| Fumarate | -1.7779 | -1.50E-15 | -6.4888 | 2.13E-05 | 0.000109 |
| Thr-Arg | -1.7722 | 4.53E-16 | -6.4 | 2.45E-05 | 0.000118 |
| Acetylaminoadipate | -1.7625 | 7.10E-16 | -6.2704 | 3.00E-05 | 0.000137 |
| NADP+ | -1.7555 | -1.51E-15 | -6.1596 | 3.58E-05 | 0.000156 |
| 3-Hydroxymethylglutarate | -1.7489 | 2.79E-15 | -6.1133 | 3.85E-05 | 0.00016 |
| CDP-ethanolamine | -1.7384 | -5.60E-16 | -5.9531 | 4.99E-05 | 0.000197 |
| alpha-Ketoglutarate | -1.7348 | 7.70E-16 | -5.8896 | 5.53E-05 | 0.000209 |
| Adenosine diphosphate | -1.6823 | -7.64E-16 | -5.3111 | 0.000146 | 0.000528 |
| Cytidine diphosphate | -1.6642 | 6.94E-16 | -5.1247 | 0.000201 | 0.000699 |
| Glycerol 3-phosphate | -1.6605 | -1.63E-15 | -5.0958 | 0.000211 | 0.000707 |
| Cys-Gly | -1.6457 | -8.22E-16 | -4.9565 | 0.00027 | 0.000866 |
| Succinate | -1.6359 | -2.34E-15 | -4.9299 | 0.000283 | 0.000866 |
| Glucose | 1.6344 | -8.95E-16 | 4.9098 | 0.000293 | 0.000866 |
| Glutathione | -1.6381 | 6.85E-16 | -4.8992 | 0.000299 | 0.000866 |
| Alanine/beta-Alanine | -1.6365 | 3.41E-15 | -4.8773 | 0.00031 | 0.000871 |
| Glu-Pro | -1.6337 | 1.81E-15 | -4.8563 | 0.000322 | 0.000876 |
| Uridine diphosphate | -1.5952 | 1.02E-16 | -4.5553 | 0.000553 | 0.001457 |
| Adenosine monophosphate | -1.5877 | -5.02E-15 | -4.5041 | 0.000607 | 0.001552 |
| Lysine | 1.5688 | -3.17E-16 | 4.4256 | 0.0007 | 0.00174 |
| Acetylacrylate | 1.5446 | -1.27E-15 | 4.2924 | 0.000894 | 0.00216 |
| Hexitols | -1.5412 | -4.26E-15 | -4.2226 | 0.001017 | 0.002391 |
| Phosphorylcholine | -1.5244 | 1.83E-16 | -4.0933 | 0.001293 | 0.00296 |
| Fructose 6-phosphate/Galactose 1-phosphate/Glucose 6-phosphate/Mannose 6-phosphate | -1.4917 | 8.86E-16 | -3.8942 | 0.001877 | 0.004188 |
| Adenosine triphosphate | -1.4813 | -1.45E-16 | -3.8583 | 0.002009 | 0.004369 |
| Cytidine monophosphate | -1.4699 | 4.79E-16 | -3.7605 | 0.002416 | 0.005127 |
| Myoinositol | -1.4479 | -1.70E-15 | -3.7374 | 0.002524 | 0.005229 |
| Glycerophosphocholine | -1.4593 | 2.44E-16 | -3.7067 | 0.002676 | 0.005413 |
| Citrate/Isocitrate | -1.4493 | 4.04E-16 | -3.6776 | 0.002828 | 0.005592 |
| Sedoheptulose 7-phosphate | -1.4178 | -4.45E-16 | -3.5334 | 0.00372 | 0.007193 |
| Glutamine | 1.3103 | -9.04E-16 | 3.1131 | 0.008317 | 0.015489 |
| Proline | -1.3291 | -1.07E-15 | -3.11 | 0.008368 | 0.015489 |
| Tyrosine | 1.3046 | -1.08E-15 | 3.034 | 0.009679 | 0.017543 |
| Betaine | 1.3048 | 2.14E-15 | 3.0228 | 0.009889 | 0.017558 |
| Octanoic Acid | 1.2858 | 1.42E-15 | 2.9775 | 0.010787 | 0.018769 |
| N-Acetylserine | -1.2663 | -1.02E-15 | -2.86 | 0.013505 | 0.023038 |
| Methionine | 1.2467 | -5.75E-16 | 2.8298 | 0.014307 | 0.023937 |
| Isoleucine/Leucine | 1.2192 | -1.88E-15 | 2.7267 | 0.017411 | 0.02858 |
| Phenylalanine | 1.2153 | 9.46E-16 | 2.7149 | 0.017807 | 0.02869 |
| Ser-Arg | -1.1843 | -1.74E-15 | -2.5806 | 0.022971 | 0.036176 |
| N-Acetylaspartate | -1.184 | -4.90E-16 | -2.5734 | 0.023286 | 0.036176 |
| Guanosine diphosphate | -1.1455 | -5.61E-16 | -2.5337 | 0.025095 | 0.038302 |

**Supplementary Table 8. Antibodies for flow cytometry.**

| **Marker** | **Fluorochrome** | **Clone** | **Company** | **Cat. Number** |
| --- | --- | --- | --- | --- |
| Glucose-Cy5 | - | - | Sigma | SML3233 |
| CAR G4S linker | AF647 | - | Cell Signaling | 69782S |
| CD8 | PerCP | RPA-T8 | Biolegend | 301030 |
| CD45RO | PE-Cy7 | UCHL1 | Invitrogen | 25-0457-42 |
| CD4 | AF700 | RPA-T4 | Biolegend | 300526 |
| CCR7 | APC/Fire750 | G043H7 | Biolegend | 353246 |
| Viability | eFluor506 | - | Invitrogen | 65-0866-14 |
| CD137 | BV605 | 4B4-1 | Biolegend | 309822 |
| CD279 | BV421 | EH12.2H7 | Biolegend | 329920 |
| CD3 | BV750 | SK7 | Biolegend | 344845 |
| IFN-γ | BV421 | 4S.B3 | Biolegend | 502508 |

**Supplementary Figure 1.** Guide RNA screening identifies a highly active gRNA at intron 2 of *ATG5*. (A) Genomic structure and target regions within intron 2 (I2) of *ATG5*. A, B, and C denote non-repetitive target DNA sequences used to design gRNAs using CRISPOR. (B) Surveyor nuclease assay showing percentage of insertions and deletions (indels) in K562 cells three days post-electroporation with SpCas9-sgRNA plasmid. Lanes labelled with “-“ represent the control amplicon for that region. Cleavage products smaller than the control amplicon represent mutations induced by editing.

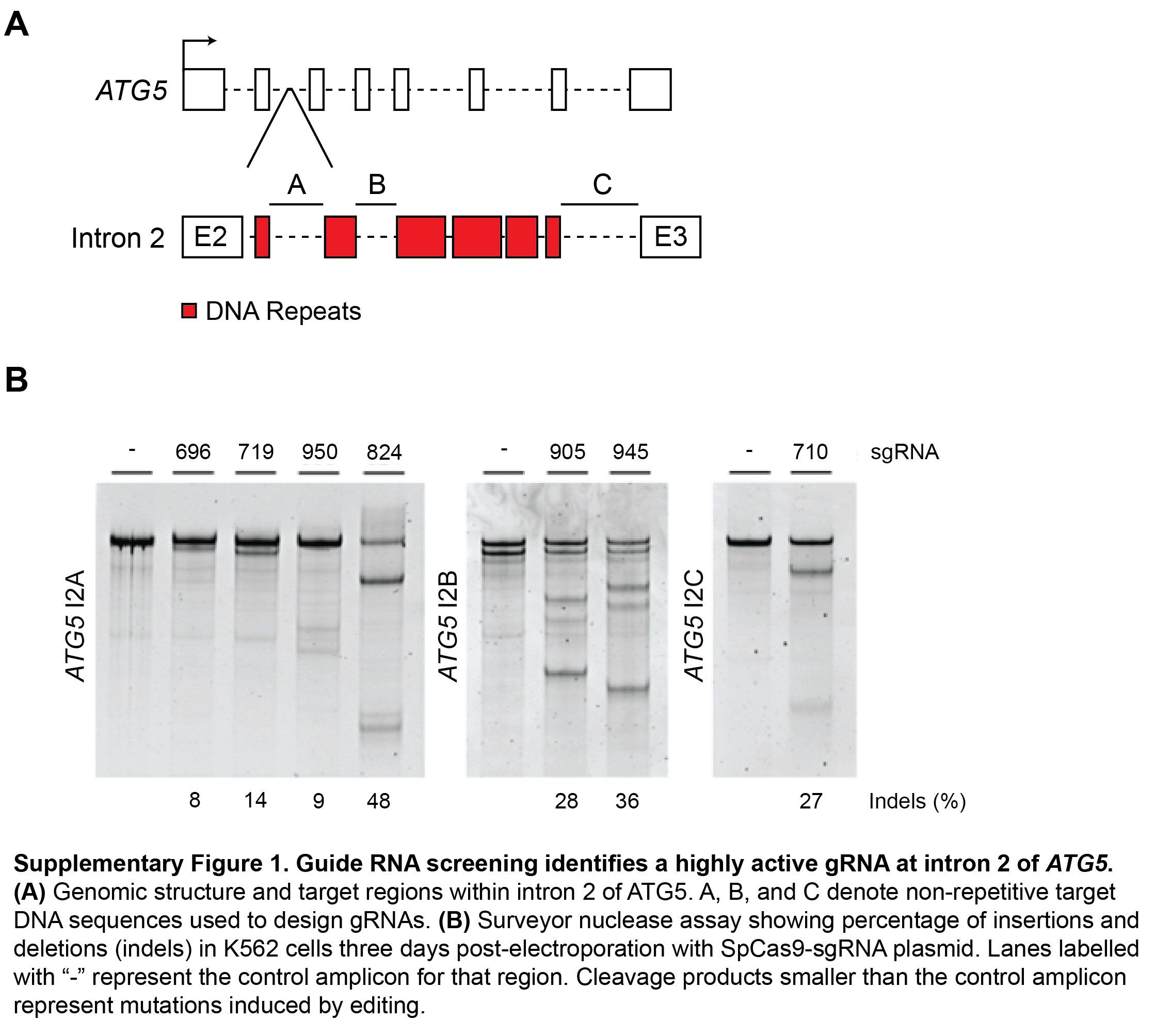

**Supplementary Figure 2.** Editing at *ATG5* eliminates functional autophagy. (A) Schematic of the plasmid donor and *ATG5* intron 2 after mScarlet-I integration. The donor contains a splice acceptor site (SA), 2A self-cleaving peptide (2A), mScarlet-I cassette, polyadenylation sequence (pA), and homology arms (HA). (B) Out-out PCR showing mScarlet-I integration at intron 2 of *ATG5*. (C) Western blot showing loss of ATG5 in K562 clones with mScarlet-I integration. (D-E) Western blots showing response to treatment with hydroxychloroquine and/or rapamycin in *ATG5*-knockout (D) or *ATG5*-complemented K562 clones (E). Loss of LC3-II indicating impaired autophagic flux is observed in clones 5, 6, 7. Restoration of autophagy function is observed in clones 5.1, 5.3, 5.6.

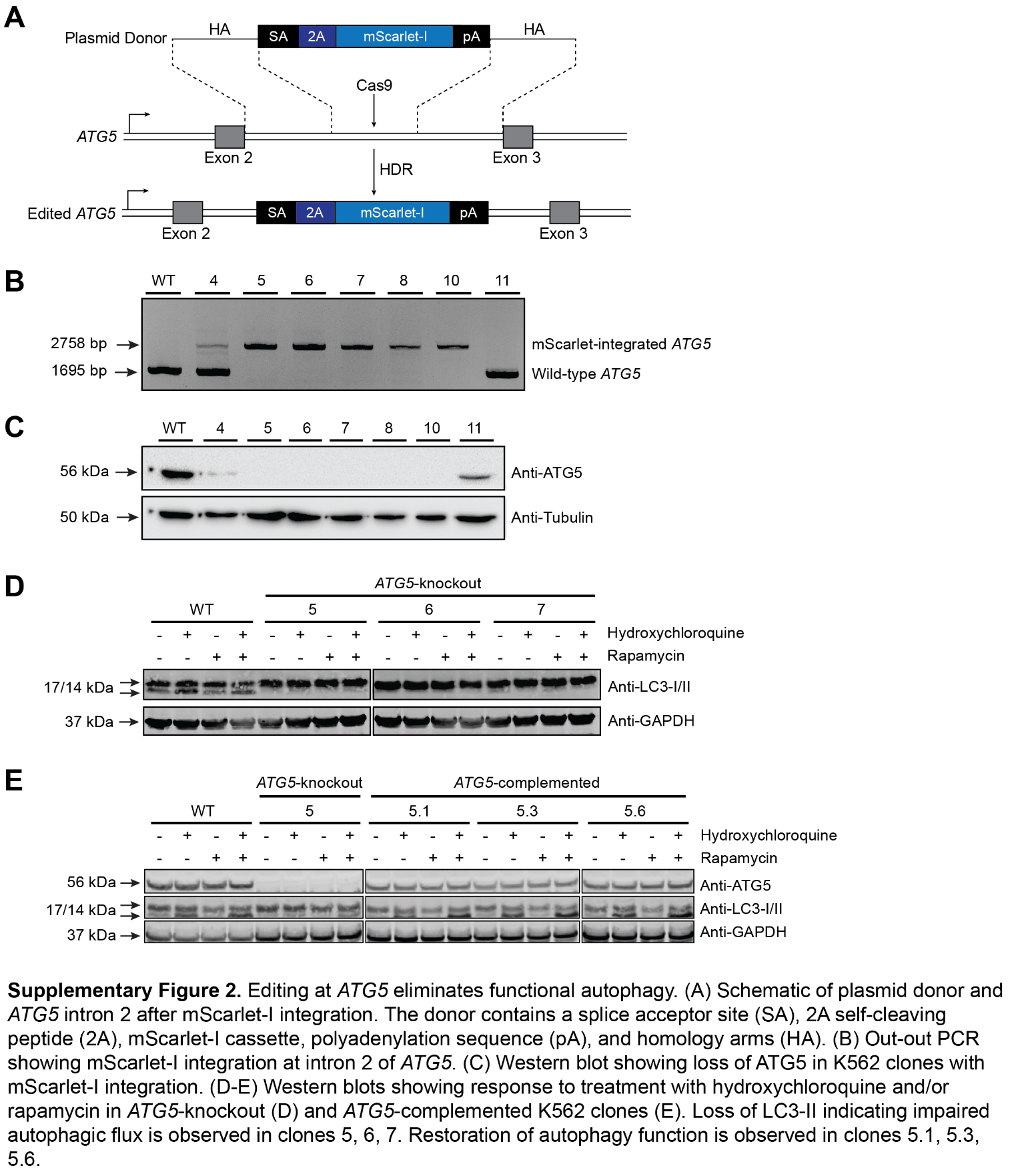

**Supplementary Figure 3.** Generating *ATG5*-knockout and *AAVS1*-knockout CAR-T cells. (A) Schematic of the AAV6 donor template and *ATG5* intron 2 after CAR integration. (B) Schematic of the AAV6 donor template and *AAVS1* intron 1 after CAR integration. Each donor contains a full-length EF1α promoter, αFR-CAR cassette, polyadenylation sequence (pA), and homology arms (HA) for the respective integration site.

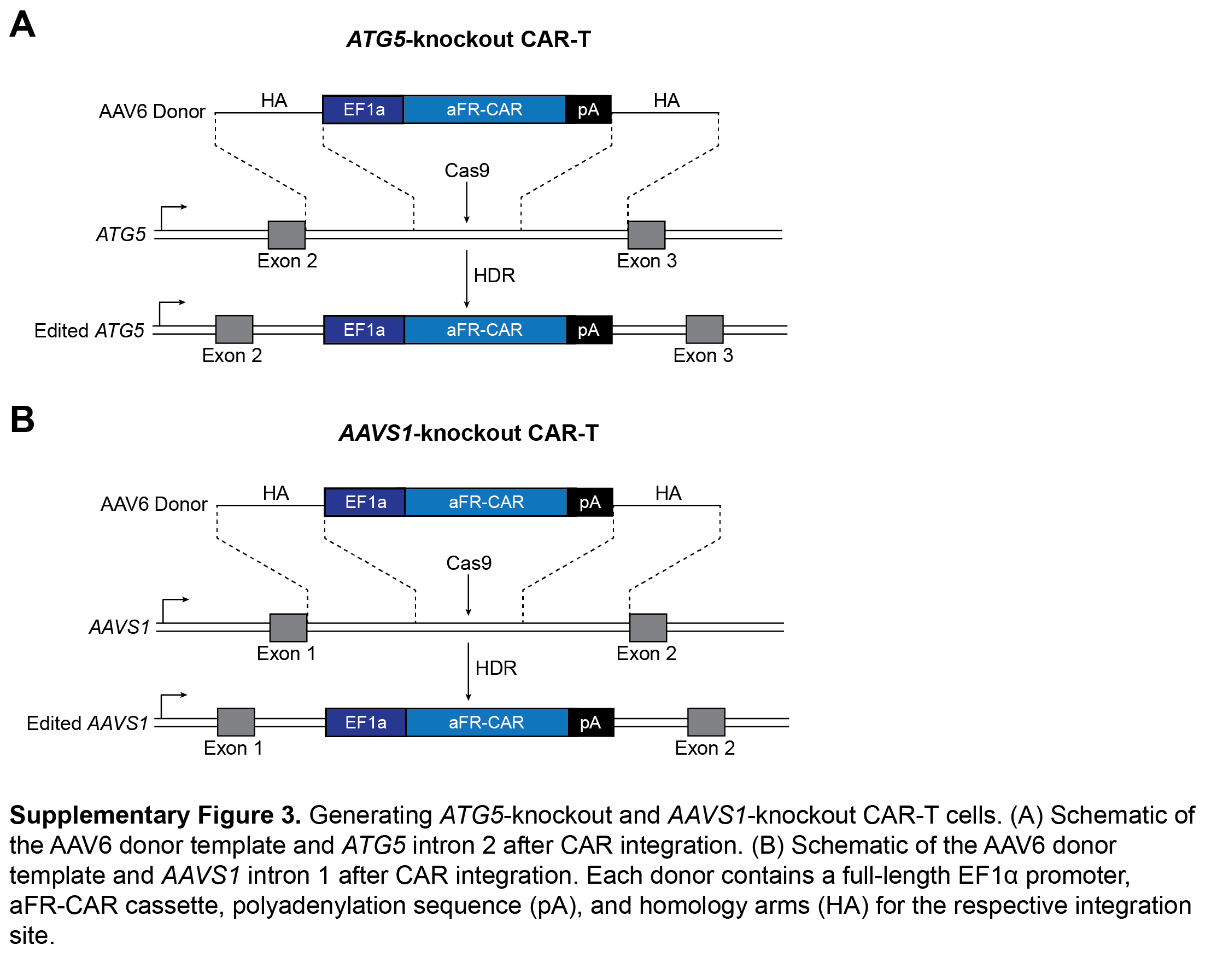
